# Temporal Variability in Brain Connectivity for Estimating Two-Year Clinical Severity in Mild Cognitive Impairment

**DOI:** 10.64898/2026.11.04.764369

**Authors:** Yury Maximov, Aleksandr Volkov, Tamoghna Chattopadhyay, Paul M. Thompson, Ildar Gabitov

## Abstract

Resting-state functional MRI measures coordinated activity across brain networks and may reveal early changes relevant to cognitive decline. Conventional ‘static’ functional connectivity analyses often compute time-series correlations across an entire scan, missing informative fluctuations over shorter timescales. We investigated whether these fluctuations, in fMRI scans collected at baseline, improve estimation of clinical severity two years later in mild cognitive impairment (MCI). In 48 ADNI participants with MCI, we summarized temporal variability using the standard deviation of windowed correlations between 21 pairs of brain networks. Using repeated nested cross-validation, the mean absolute error in predicting follow-up Clinical Dementia Rating Sum of Boxes (CDR-SB) was 1.62 points with clinical covariates alone, 1.66 after adding static connectivity, and 1.51 after adding connectivity variability. With additional motion covariates, the corresponding errors were 1.70, 1.73, and 1.59. Connectivity variability outperformed both conventional models in all three cross-validation repetitions, while randomized signals preserving the complete cross-spectrum did not reproduce the improvement. Perhaps surprisingly though, most of the gain occurred in participants without subsequent clinical worsening, and results were sensitive to the choice of temporal windows. These preliminary findings suggest that compact measures of connectivity variability may capture clinically relevant information beyond conventional measures of ‘static’ functional connectivity, motivating larger studies to evaluate their value for predicting future cognitive impairment.

## 1 Introduction

People with mild cognitive impairment (MCI) vary considerably in their risk of future cognitive decline and dementia. Estimating their future clinical severity is important for patient monitoring and selecting participants for clinical trials. Resting-state functional MRI offers a noninvasive approach to study coordinated activity across brain networks, revealing differences that are not apparent from clinical assessments alone [1]. Conventional ‘static’ functional connectivity (FC) measures summarize correlations between brain regions over an entire scan. However, two individuals with similar average connectivity may differ substantially in how their connections fluctuate over shorter periods. We investigate whether these fluctuations provide additional information relevant to predicting clinical impairment two years later, beyond baseline clinical status and static FC.

Prior studies have used static and dynamic FC to characterize MCI and predict conversion to Alzheimer’s disease [2, 3, 4, 5, 6, 7]. Dautricourt et al. found that dynamic connectivity states were associated with dementia risk factors, including vascular health and cognitive activity, in cognitively normal older adults [4]. Hu et al. showed that switching between functional network configurations was associated with cognitive performance in MCI, with reduced default mode network switching predicting progression to dementia [5]. Shakil et al. demonstrated that sliding-window connectivity estimates are sensitive to window length and spacing, motivating careful control of temporal sampling when interpreting connectivity fluctuations [8].

Here, we focus on predicting the continuous Clinical Dementia Rating Sum of Boxes (CDR-SB) score, a common measure of the severity of cognitive and functional impairment, often used in clinical trials. We summarize connectivity variability using just 21 features, representing the standard deviation of correlations across time windows for each pair of seven resting-state networks. This compact representation preserves the pattern of variability across brain networks, which may contain clinically relevant information missed by conventional static FC.

A key challenge is that apparent connectivity variability depends on scan duration, motion, and how temporal windows are selected [8, 9, 10, 11]. We compare clinical predictors, static FC, connectivity variability, and their combination using the same participants and cross-validation procedures. We test whether variability adds information beyond clinical and static connectivity measures, and examine individual prediction errors, spectral controls, and alternative temporal sampling strategies to investigate sources of improvement. These exploratory analyses assess whether compact measures of network variability offer a promising approach to estimate future clinical impairment and warrant larger independent studies.

## 2 Data and Methods

### 2.1 Participants and Clinical Outcome

To evaluate whether resting-state connectivity variability helps estimate future clinical severity, we analyzed data from the Alzheimer’s Disease Neuroimaging Initiative (ADNI) [12]. The initial dataset contained 4,347 Blood Oxygenation Level Dependent (BOLD) fMRI series from 1,764 participants. Clinical and imaging eligibility criteria, including diagnosis within 90 days of MRI and adequate scan duration, retained 1,698 participants. From those meeting additional longitudinal and imaging requirements, we matched 90 participants with dementia to 90 cognitively normal (CN) participants and 90 with mild cognitive impairment (MCI), using visit count, demographic characteristics, and acquisition parameters. The resulting cohort comprised 270 participants and 732 fMRI visits.

Our prediction analysis focused on 48 participants with MCI who had baseline and approximately two-year follow-up Clinical Dementia Rating Sum of Boxes (CDR-SB) assessments [13]. Baseline CDR-SB was measured within 90 days before MRI, and follow-up was selected closest to two years. The outcome was future CDR-SB severity, rather than change from baseline or conversion to dementia.

The clinical model included baseline CDR-SB, age, sex, repetition time (TR), scan length, mean framewise displacement, scanner, acquisition phase, and days to the future assessment. Prediction was retrospective and conditional on observed follow-up time. An additional motion analysis added the fraction of frames exceeding 0.2 mm displacement and median standardized DVARS to every model. All improvements were calculated between models with the same covariate set.

### 2.2 MRI Representations

To reduce motion-related effects, we jointly regressed out an intercept, linear trend, and 24 motion parameters (M24). BOLD time series from the Schaefer-400 parcels [14] were then averaged within the seven Yeo resting-state networks [15].

We compared two representations of functional connectivity (FC). **Static FC** measured the Pearson correlation between each pair of networks across the entire scan, giving 21 features. To measure how these correlations varied over time, we also calculated FC in successive, overlapping windows of approximately 60 seconds, advanced every 30 seconds. Window length w and step size h were adjusted for each scan’s repetition time (TR): *w* = max(8, [60*/*TR]) and *h* = max(1, [30*/*TR]), where [·] is a rounding operator. For each network pair e, we quantified connectivity variability (dispersion) as the standard deviation of its correlations across *K* windows:

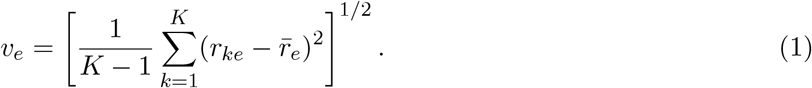

Here, *r*_*ke*_ is the correlation for network pair e in window k, and 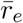 is its mean across windows. Correlations were not Fisher-transformed. The resulting 21 dispersion features capture which network connections vary most, without retaining the order of the windows. We evaluated static FC (s), dispersion (v), and their combination [s,v], comprising 42 features, to test whether temporal variability adds predictive information beyond average connectivity.

### 2.3 Model Fitting and Evaluation

To assess whether connectivity variability improves prediction of future clinical severity, we compared models using the same participants and cross-validation procedure. Given the small sample (N = 48), we used five-fold participant-level cross-validation, repeated three times with different random splits. Within each training set, three-fold cross-validation selected the number of partial least squares (PLS) components (2, 5, or 10) and ridge regularization strength (0.1, 1, 10, or 100). This allowed model complexity to be optimized without using test data [16]. MRI features were standardized using training data and scaled by the square root of the feature-block dimension to balance their contributions. PLS used the training outcomes without additional outcome scaling. Clinical covariates and PLS scores were also standardized within training folds, ensuring that all preprocessing and model selection excluded held-out participants.

Our primary comparison tested whether adding connectivity dispersion to clinical covariates and static FC reduced prediction error. We also compared dispersion directly with clinical-only and static-FC models. Performance was measured by mean absolute error (MAE) in follow-up CDR-SB points, averaged across 3 cross-validation repetitions. As these repetitions used the same participants, they assess sensitivity to data partitioning rather than providing independent replication.

## 3 Results

### 3.1 Model Fitting and Evaluations

We tested whether connectivity variability improves estimation of 2-year clinical impairment by comparing clinical, static FC, dispersion, and combined models in the same 48 participants. To limit overfitting, we used nested cross-validation, with 5 outer folds for evaluating prediction accuracy and 3 inner folds to select model parameters. This was repeated 3 times with random splits. The inner folds selected 2, 5, or 10 partial least squares (PLS) components and a ridge penalty of 0.1, 1, 10, or 100. The clinical-only model used the same inner folds to select its penalty.

MRI features were standardized using training data and scaled by the square root of the number of features to balance their contributions. Clinical covariates and PLS scores were similarly standardized. All transformations and model selection were restricted to training data, keeping test participants independent of model fitting.

Our primary test was whether adding dispersion to clinical covariates and static FC reduced prediction error. We also compared dispersion with the clinical-only and static-FC models. Accuracy was measured by mean absolute error (MAE) in follow-up CDR-SB points, averaged across the three repetitions. These repetitions assess sensitivity to data splitting rather than independent replication. As this was an exploratory study, the findings require validation in independent participants.

To assess whether the prediction gain could arise by chance during model selection, we repeated the analysis after randomly shuffling MRI features across participants 99 times, preserving relationships among MRI feature groups. A gain at least as large as observed occurred once for the selected model, but five times when selecting the best of 24 candidate models. As shuffling disrupts relationships between MRI features and clinical covariates, this exploratory control is not a formal significance test.

### 3.2 Participant Contributions to the Mean Gain

We examined individual prediction errors to determine whether the improvement was consistent across participants.

Connectivity dispersion reduced absolute error in 24 of 48 participants, or 25 when additional motion covariates were included (Fig. 2). Among 29 participants whose CDR-SB scores worsened, mean improvements were small (0.02 and −0.01 points, respectively). The larger gains occurred among 19 participants without worsening (0.26 and 0.31 points). These groups were identified from clinical outcomes and were not used in model training.

**Figure 1.**
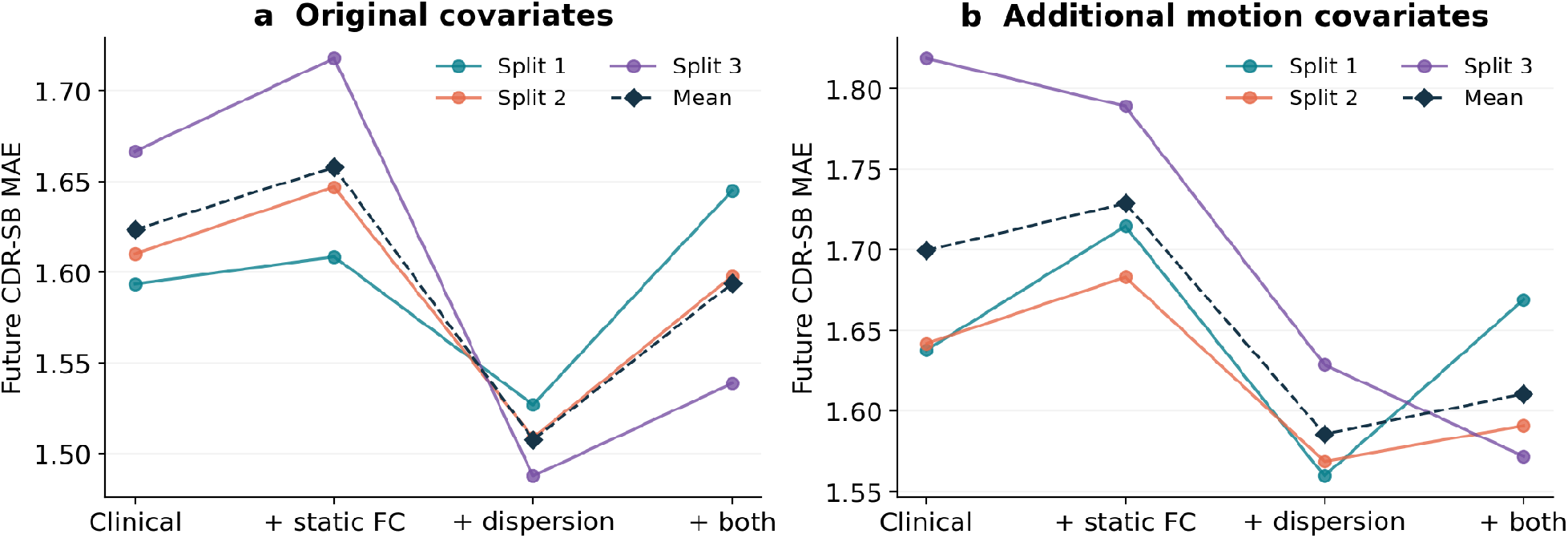
Prediction of two-year CDR-SB severity in 48 participants with MCI. Mean absolute errors (MAE) are shown for clinical covariates alone and with static FC, connectivity dispersion, or both. Panels compare original (a) and additional motion covariates (b). Lines show three cross-validation repetitions; diamonds indicate means. Dispersion alone achieved the lowest mean error in both analyses.

**Figure 2.**
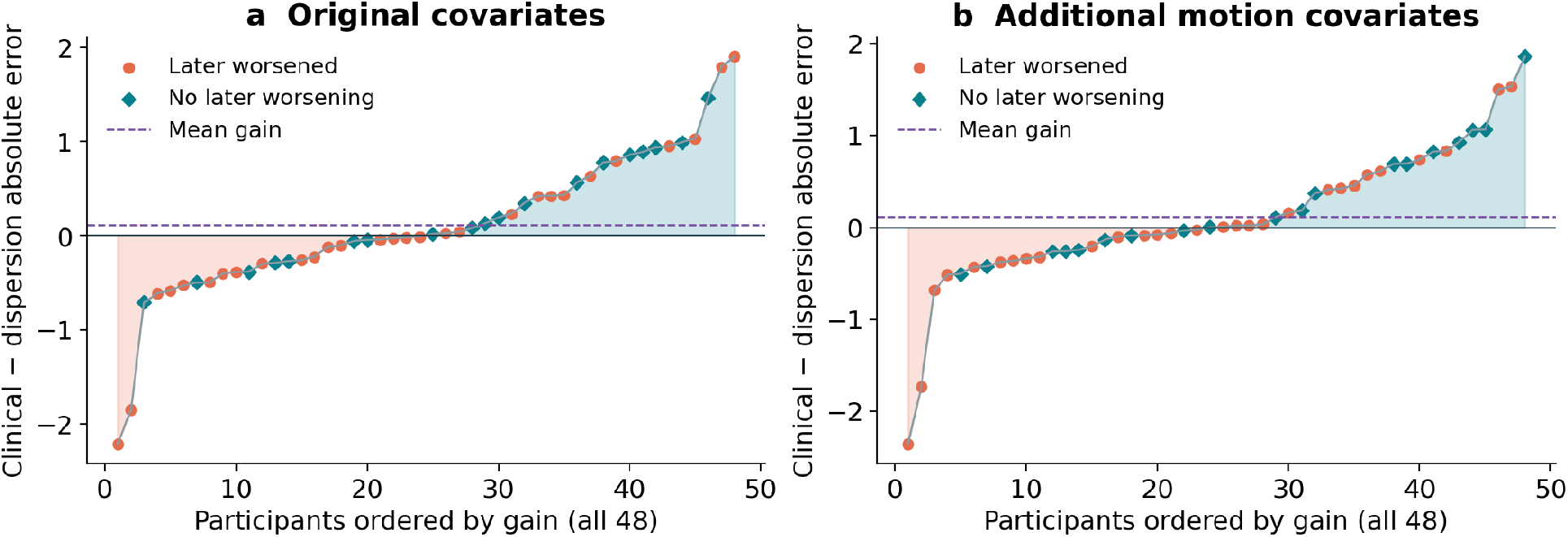
Improvement in individual prediction accuracy after adding connectivity dispersion to clinical covariates. Positive values indicate lower absolute CDR-SB prediction error. Each point represents one participant, averaged across three cross-validation repetitions. Symbols distinguish participants with and without subsequent clinical worsening; dashed lines indicate mean improvement. Participants are ordered separately by improvement in each panel.

The average benefit remained positive when any single participant’s contribution was removed, but disappeared when the five largest contributions were excluded. Improvements also differed between ADNI2 and ADNI3 (0.27 versus −0.02 points, or 0.22 versus 0.02 with additional motion covariates). These findings suggest that connectivity variability may improve estimates of future clinical severity, although the benefit is not uniform across participants or acquisition cohorts.

To test whether differences in the timing of clinical assessments affected our results, we repeated the analysis in 38 participants whose baseline CDR-SB was measured within 30 days before MRI. After retraining the models, clinical-only and dispersion MAEs were 1.70 and 1.60 points, respectively, or 1.76 and 1.63 with additional motion covariates. Dispersion improved accuracy in two of three cross-validation repetitions, and all three with additional motion covariates. Thus, the average benefit persisted when baseline assessments were closer to MRI.

### 3.3 Contribution of Individual Network Pairs

We next investigated which brain network connections contributed most to prediction accuracy. We removed each of the 21 dispersion features in turn and retrained the models. Excluding Limbic–Visual, Limbic–Salience/Ventral Attention, or Salience/Ventral Attention–Somatomotor connectivity increased MAE by 0.14, 0.10, and 0.09 points, respectively, in all three repetitions. Removing some other connections improved accuracy. These findings suggest that variability in specific network connections may be particularly informative, but correlated features prevent us from attributing the effects to individual connections.

## 4 Spectral and Temporal Controls

**Table 1.** Future CDR-SB MAE in points. Every model includes the corresponding clinical and technical covariates.

| MRI representation | Original | + motion |
|---|---|---|
| None | 1.623 | 1.699 |
| Static FC | 1.658 | 1.729 |
| Observed dispersion | 1.508 | 1.586 |
| Static + dispersion | 1.594 | 1.611 |
| Mean spectral reference | 1.703 | 1.825 |
| Observed minus reference | 1.518 | 1.613 |
| Reference + residual | 1.681 | 1.731 |

### 4.1 Comparison with Spectral-Reference Features

An important question is whether connectivity variability provides information beyond the frequency spectra and correlations already present in conventional fMRI measurements. To test this, we generated 39 randomized reference signals for each scan by applying the same random Fourier phase to all seven networks at each frequency. This preserves the complete cross-spectrum and overall functional connectivity while changing the temporal patterns of the signals [17].

We compared the observed dispersion features (v) with their mean across randomized references (m), their difference (v−m), and the combined features [m, v −m]. Using identical participants and cross-validation splits, we tested whether the observed variability provides predictive information beyond that reproduced by the spectral properties of the signals.

The randomized spectral references performed worse than the clinical-only model in every cross-validation repetition, suggesting that conventional spectral properties alone did not reproduce the predictive benefit of observed connectivity variability. Subtracting the mean reference dispersion from the observed features retained improvements of 0.11 and 0.09 CDR-SB points, without and with additional motion correction, respectively. Combining the reference and residual features did not improve accuracy further. Results also depended on which randomized references were used: two independent sets gave MAEs of 1.80 and 1.71, indicating some sensitivity to surrogate sampling. Overall, the observed connectivity fluctuations contained predictive information not reproduced by the spectral controls.

### 4.2 Effects of Window Selection and Temporal Coverage

We next examined whether prediction accuracy depended on which parts of the scan were used to estimate connectivity variability. Using eight consecutive windows from the beginning, middle, or end of each scan gave MAEs of 1.64–1.65, whereas eight windows distributed across the scan improved MAE to 1.56, approaching the full-scan result of 1.51. Sampling a longer period may provide more informative estimates of connectivity variability.

However, distributed windows may also cover more distinct time points. To separate these effects, we repeated the comparison using four nonoverlapping windows, so that each selection contained exactly the same number of time points. Under this stricter control, distributed windows performed worse (MAE = 1.73) than consecutive windows (1.63–1.67), with similar findings after additional motion adjustment (Fig. 3). Thus, the apparent benefit of distributing windows across a scan depends on the sampling strategy, highlighting the importance of controlling temporal coverage when developing connectivity-based biomarkers.

**Figure 3.**
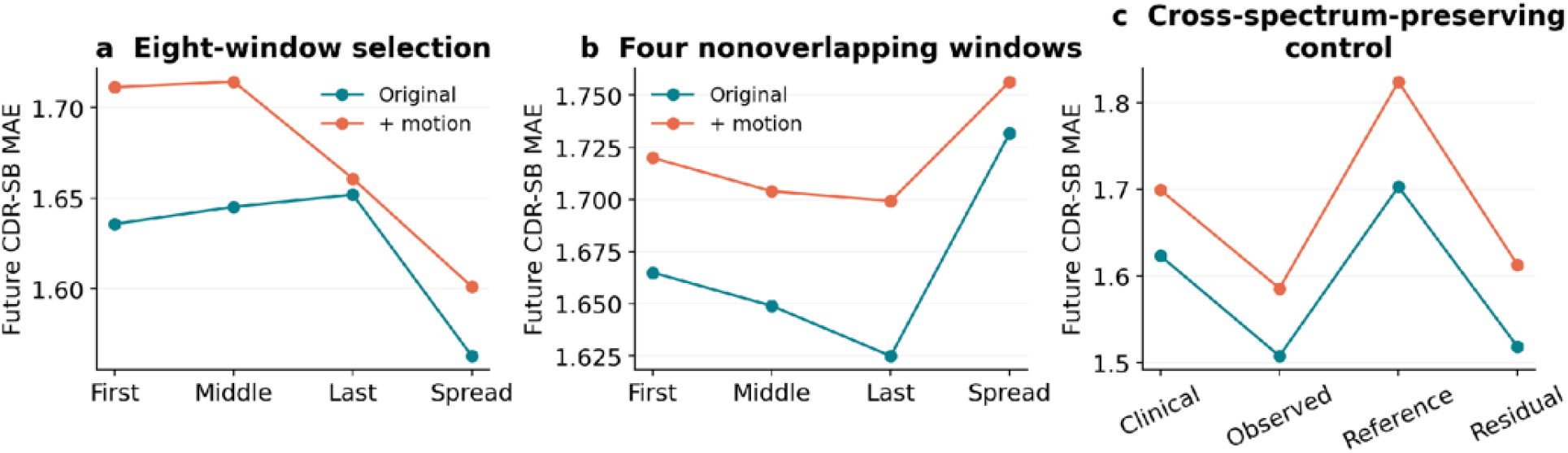
Effects of temporal sampling and spectral controls on two-year CDR-SB prediction. (a) Eight windows distributed across the scan outperform consecutive windows. (b) This advantage disappears when four nonoverlapping windows contain equal numbers of time points. (c) Dispersion from randomized signals preserving the cross-spectrum does not reproduce the predictive gain, whereas the observed-minus-reference features retain it. Values are mean errors across three cross-validation repetitions. Window count alone does not determine dependence among correlation estimates. Overlap, spacing, and temporal coverage also affect that dependence. Nuisance regression was fitted to the full record in all of these retrospective comparisons. The advantage for eight distributed windows did not extend to the four-window design.

## 5 Discussion and Conclusion

This pilot study suggests that temporal variability in brain connectivity may provide information about future cognitive impairment beyond conventional static functional connectivity (FC). A compact set of just 21 connectivity-dispersion features improved estimation of two-year CDR-SB severity compared with clinical and static-FC models. The improvement persisted after additional motion adjustment and stricter clinical assessment timing. Randomized signals preserving the complete cross-spectrum did not reproduce the predictive gain, suggesting that observed connectivity variability captures additional temporal information. These findings extend previous studies of dynamic connectivity in MCI [2, 3] by examining continuous clinical severity and investigating the spectral and temporal origins of predictive information.

Several limitations remain. Most of the improvement occurred in participants without subsequent clinical worsening. Results also depended on temporal window selection, and variability in windowed correlations does not necessarily indicate changes in neuronal connectivity [18, 19, 20]. The sample was small (N = 48), and the exploratory analyses require independent validation [21]. Nevertheless, these preliminary results suggest that compact measures of connectivity variability may complement conventional fMRI features in estimating future clinical impairment. Larger longitudinal studies should test their generalizability and stability [22], optimize temporal sampling, and examine their ability to predict cognitive decline. This approach offers a promising direction for developing functional MRI biomarkers of dementia progression.

## 6 Code and Data Availability

Prediction code, a reproducible synthetic example, and aggregate results from the 48-participant ADNI analysis are available at https://github.com/yury-rd/mci-forecast.

## 7 Compliance with Ethical Standards

We analyzed previously collected, publicly available ADNI data, without recruiting participants or acquiring new scans. For this analysis of public, de-identified anonymized data, no additional institutional review board approval was required.

## Acknowledgments

Data used in preparation of this article were obtained from the Alzheimer’s Disease Neuroimaging Initiative (ADNI) database (adni.loni.usc.edu). As such, the investigators within the ADNI contributed to the design and implementation of ADNI and/or provided data but did not participate in the analysis or writing of this report. A complete listing of ADNI investigators can be found in the ADNI investigator acknowledgment list.

The ADNI imaging and clinical data came from the Alzheimer’s Disease Neuroimaging Initiative database. ADNI began in 2003 as a public-private partnership led by Michael W. Weiner, MD. It brings together longitudinal imaging, biomarkers, and clinical and neuropsychological assessments to study progression from mild cognitive impairment to Alzheimer’s disease.

The National Institute on Aging funds ADNI data collection and sharing through National Institutes of Health grant U19 AG024904 to the Northern California Institute for Research and Education. ADNI has also received support from the National Institute of Biomedical Imaging and Bioengineering, the Canadian Institutes of Health Research, and private contributors through the Foundation for the National Institutes of Health. These contributors include AbbVie, Alzheimer’s Association, Alzheimer’s Drug Discovery Foundation, Araclon Biotech, BioClinica, Inc., Biogen, Bristol-Myers Squibb Company, CereSpir, Inc., Cogstate, Eisai Inc., Elan Pharmaceuticals, Inc., Eli Lilly and Company, EuroImmun, F. Hoffmann-La Roche Ltd and its affiliated company Genentech, Inc., Fujirebio, GE Healthcare, IXICO Ltd., Janssen Alzheimer Immunotherapy Research & Development, LLC., Johnson & Johnson Pharmaceutical Research & Development LLC., Lumosity, Lundbeck, Merck & Co., Inc., Meso Scale Diagnostics, LLC., NeuroRx Research, Neurotrack Technologies, Novartis Pharmaceuticals Corporation, Pfizer Inc., Piramal Imaging, Servier, Takeda Pharmaceutical Company, and Transition Therapeutics.

